# Modulation of prion protein expression through cryptic splice site manipulation

**DOI:** 10.1101/2023.12.19.572439

**Authors:** Juliana E. Gentile, Taylor L. Corridon, Meredith A. Mortberg, Elston Neil D’Souza, Nicola Whiffin, Eric Vallabh Minikel, Sonia M. Vallabh

## Abstract

Lowering expression of prion protein (PrP) is a well-validated therapeutic strategy in prion disease, but additional modalities are urgently needed. In other diseases, small molecules have proven capable of modulating pre-mRNA splicing, sometimes by forcing inclusion of cryptic exons that reduce gene expression. Here, we characterize a cryptic exon located in human *PRNP*’s sole intron and evaluate its potential to reduce PrP expression through incorporation into the 5’ untranslated region (5’UTR). This exon is homologous to exon 2 in non-primate species, but contains a start codon that would yield an upstream open reading frame (uORF) with a stop codon prior to a splice site if included in *PRNP* mRNA, potentially downregulating PrP expression through translational repression or nonsense-mediated decay. We establish a minigene transfection system and test a panel of splice site alterations, identifying mutants that reduce PrP expression by as much as 78%. Our findings nominate a new therapeutic target for lowering PrP.

## Introduction

Prion disease is a rapidly fatal neurodegenerative disease caused by the templated misfolding of the prion protein, PrP, encoded by the prion protein gene (*PRNP* in humans)^1^. Prion disease naturally afflicts a range of mammals and has long been modeled in laboratory rodents, in which the full disease process can be induced. Both genetic^2^ and pharmacological^3,4^ experiments in such models have demonstrated that reducing the amount of PrP in the brain is protective against prion disease, inspiring hope that a PrP-lowering therapy could be used to effectively treat, delay, and prevent disease in patients and individuals at risk^5^. An RNAse H1 antisense oligonucleotide targeting *PRNP* RNA for degradation is now in preclinical development^3,4,6,7^, but additional therapeutic candidates are urgently needed.

Recently, the FDA-approved drug risdiplam^8–11^ and clinical candidates kinetin and branaplam^12–15^ have highlighted small molecule modulation of pre-mRNA splicing as another tool for therapeutic tuning of gene expression. Branaplam causes incorporation of a piece of intronic sequence — variously called a non-annotated exon, cryptic exon, or poison exon — into mature *HTT* mRNA, causing a frameshift and nonsense-mediated decay^15^. Inspired by this work, we were led to inquire whether the architecture of *PRNP* would lend itself to disruption via splice site manipulation. *PRNP*’s coding sequence is located entirely within a single exon, precluding frameshift strategies. We hypothesized, however, that inclusion of a novel upstream open reading frame (uORF) in the *PRNP* 5’UTR could decrease PrP expression. It is known that uORFs can have dramatic effects on gene expression^16,17^ either through reduced abundance of ribosomes on the canonical ORF, or possibly through nonsense-mediated decay (NMD) triggered by the presence of a stop codon prior to the final splice junction, though the latter mechanism is debated^18^. The existence of Mendelian diseases caused by variants introducing uORFs^19^, the evolutionary constraint of genetic variants that cause or extend uORFs in dosage sensitive genes^20^, as well as work with uORF-targeting antisense oligonucleotides^21^, underscore the potential functional impact of uORFs.

Here, we identified a potential uORF within a cryptic exon located in *PRNP*’s sole intron, homologous to exon 2 in many non-primate species. By genetically strengthening the splice sites surrounding the cryptic exon located in *PRNP*’s 5’ UTR, we show that the mutations yielding the most robust inclusion of exon 2 reduced PrP expression by up to 78% in human cells. Certain other mutants reduced *PRNP* transcript levels and PrP protein expression without yielding cryptic exon inclusion detectable by qPCR, suggesting multiple mechanisms may be operative. These efforts nominate a novel strategy for lowering PrP.

## Results

*PRNP* is a small gene of roughly 15 kilobases (kb) in humans (Figure 1A). In all mammals, the entire coding sequence is contained in the final exon of the gene, while the 5’UTR is divided across exons, however, the number of exons differs. In mouse and most other preclinical species of interest, there are 3 constitutive exons^22,23^, with introns 1 and 2 dividing the 5’UTR (Figure 1B). In Syrian hamsters, exon 2 is subject to variable splicing and is included in ∼27% of transcripts^24^ (Figure 1B). In humans and several closely related primate species, *PRNP* has only two annotated exons, the equivalent of exons 1 and 3 from other mammals; exon 2 remains as a cryptic exon within the sole intron^25^. For clarity, herein we will refer to human *PRNP* exons 1, 2, and 3, and introns 1 and 2, even though the naturally occurring *PRNP* transcript contains only 2 exons and 1 intron.

**Figure 1.**
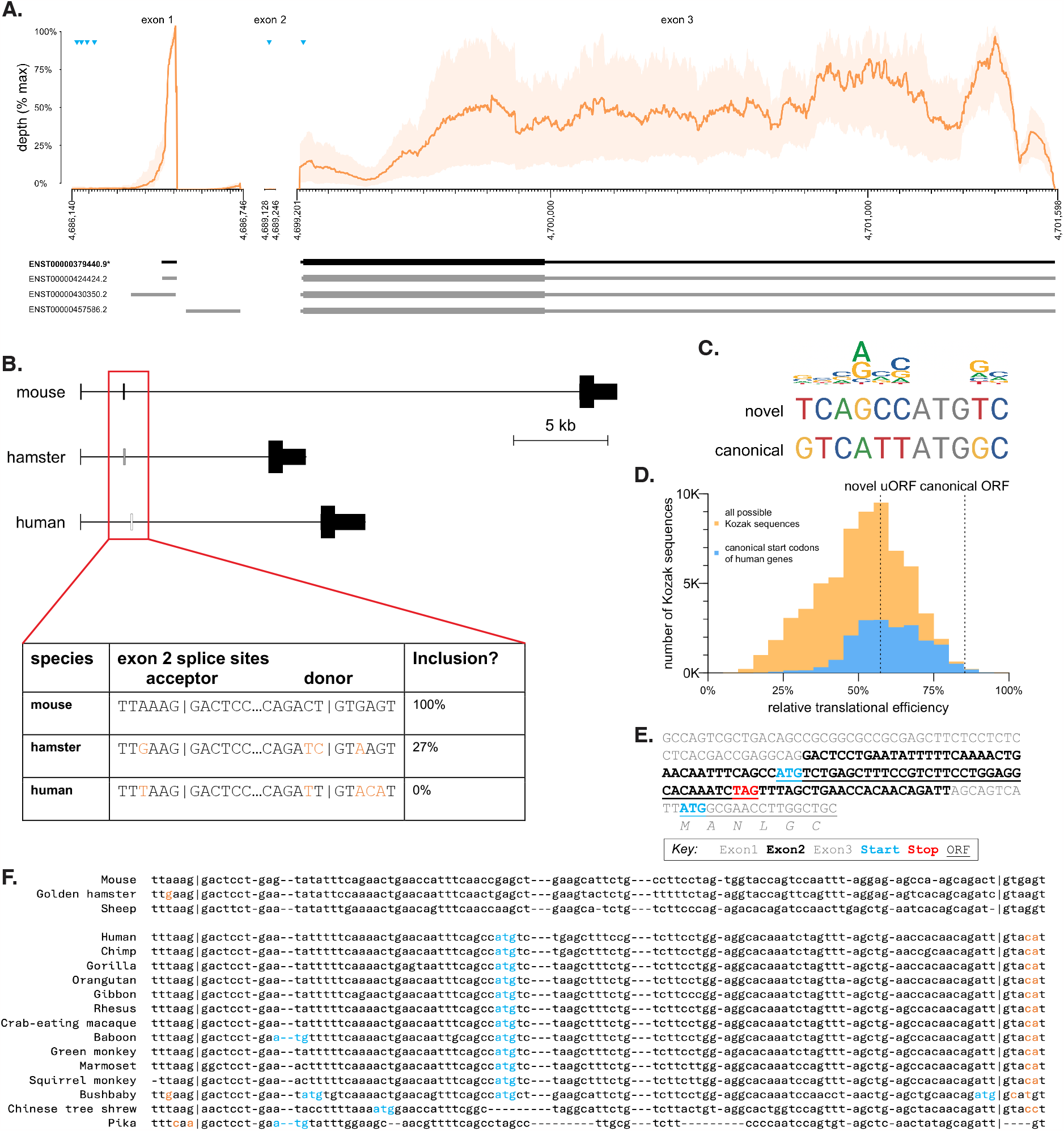
A cryptic exon in human PRNP. **A)** Human PRNP transcript structure in human brain. Top panels show GTEx^29^ v8 bulk RNA-seq coverage — mean (orange lines) and range (orange shaded area) across 13 brain regions. Coverage depth for exons 1 and 2 is normalized to the max for exon 1; depth for exon 3 is normalized to the max for exon 3. ATGs representing candidate upstream open reading frames and the canonical open reading frame are shown as blue triangles. Ensembl GRCh38.p14 annotated transcripts are shown below, canonical in black, alternatives in gray. **B)** Comparison of orthologous exon 2 sequence in mouse, hamster, and human. Hamster inclusion percentage from ref ^24^. **C)** Comparison of PRNP canonical and exon 2 novel ATG Kozak contexts with a sequence logo of human initiation sites (see Methods). **D)** Relative strength of canonical and novel PRNP ORFs in context. Shown for comparison are histograms of translational efficiency of all 65,536 (4^8) possible Kozak contexts (yellow) and of all 18,784 actual human canonical ORF Kozak contexts (blue), expressed as a percentage of the translation of the most efficient Kozak context, TTCATC<u>ATG</u>CA, according to data from Noderer et al^27^. **E)** Annotated sequence of the PRNP 5’UTR if exon 2 were included. Frame is relative to the canonical ORF, and percentile indicates strength of the Kozak context as a percentile of all possible Kozak sequences, using rankings from ref ^27^. **F)** Multiple alignment of PRNP exon 2 sequences known to be constitutively or variably included in mRNA from mouse^22^, hamster^24^, and sheep^23^ versus all orthologous sequences in eutherian mammals that contain ATGs. ATGs are shown in blue, splice site variants absent from mouse, hamster, or sheep are shown in orange. A full alignment including all eutherian mammals is shown in Figure S1.

Although essential splice sites — AG at the A-1 and A-2 and GT at the D+1 and D+2 positions — are conserved in human exon 2, we hypothesized that other nearby base pair substitutions may contribute to exclusion of this exon, particularly the loss of the G at the highly constrained D+5 position^26^ (Figure 1B). Human *PRNP* exon 2 contains an ATG in a moderately strong Kozak context (Figure 1C), estimated to yield 57% maximal translational efficiency, near the median of canonical ORFs of all other human protein-coding genes^27^ (Figure 1D). Human *PRNP* was previously reported^28^ to already contain 4 uORFs in exon 1, however, RNA-seq data from human brain tissue^29^ provide no support for transcription initiation beginning this far upstream: mean RNA-seq coverage at these uORFs is <0.5% of the peak coverage within exon 1 (Figure 1A). Thus, if exon 2 were included, its ATG would yield a new, sole uORF upstream of *PRNP*’s canonical start codon (Figure 1E) with the potential to downregulate PrP expression through its impact on ribosomal activity^16,30^. Its stop codon also occurs 22 bp prior to the exon 2/3 splice junction (Figure 1E), creating a possible opportunity to trigger nonsense-mediated decay (NMD; see Discussion). Alignment of *PRNP* exon 2 sequences across all available mammalian species (Figure 1F and Figure S1) reveals that exon 2 ATGs are present only in species with exon 2 splice site variants known or predicted to exclude exon 2 from mature mRNA, consistent with the possibility of exon 2 uORF having a strong negative effect on PrP expression. We thereby hypothesized that acting through either of these mechanisms, inclusion of exon 2 and thus the uORF of interest in *PRNP* mRNA would reduce PrP expression.

To test this hypothesis, we first sought to generate a *PRNP* minigene system to support facile splice site manipulation, transfection, and screening in cell culture. A 4.8 kb minigene lacking most of intron 1 yielded no detectable PrP expression in HEK293 cells by Western blot (Figure S2). A 6.5 kb construct retaining all of intron 1 and only the first and last 500 bp of intron 2 (Figure 2A) expressed robustly, and was used for all subsequent experiments. Codon optimization of exon 3 allowed for qPCR primer/probe pairs to discriminate minigene *PRNP* RNA from endogenous *PRNP* RNA (Figure 2B).

**Figure 2.**
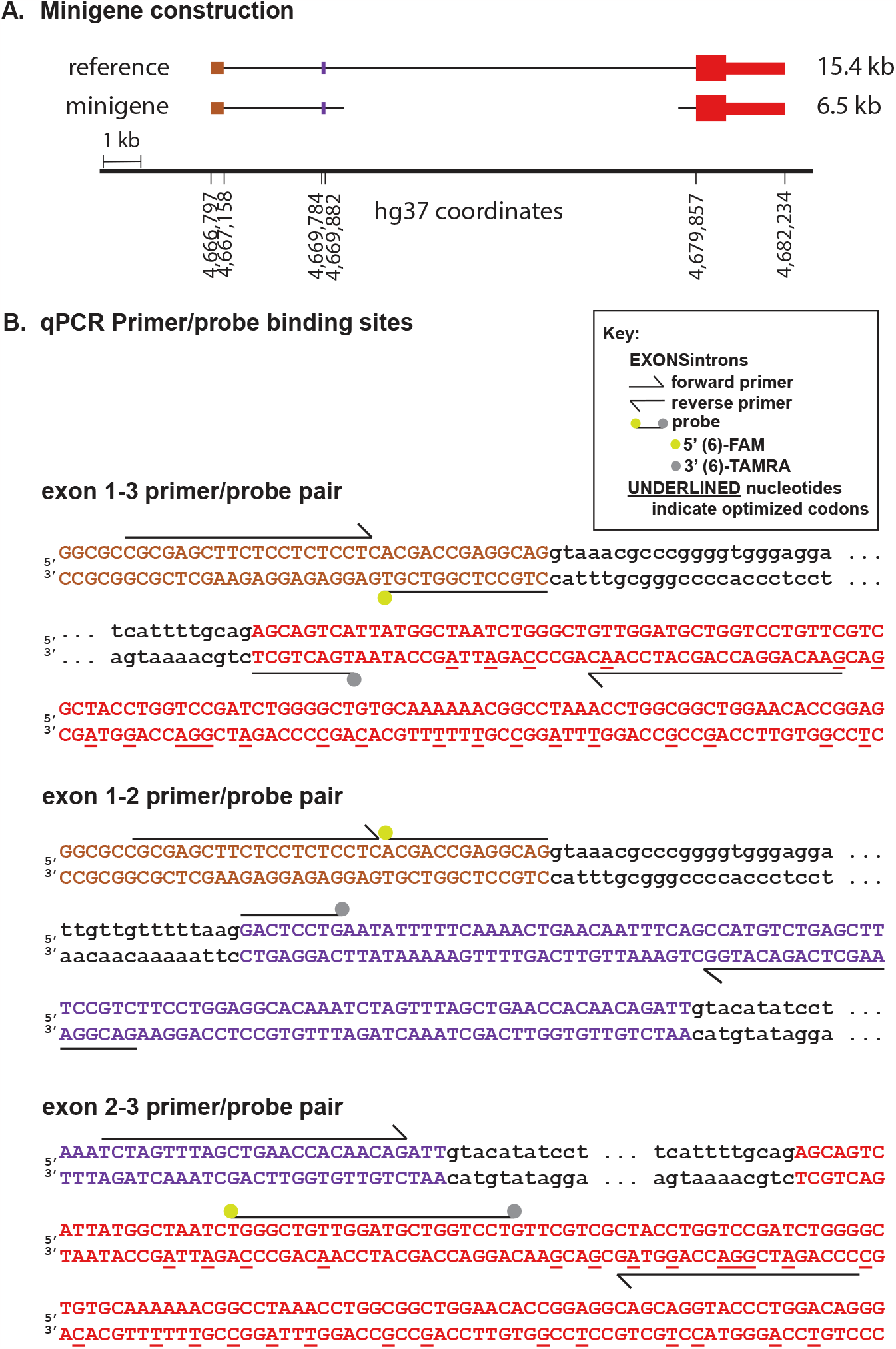
Design of PRNP minigene and primer pairs. **A)** Diagram of minigene versus human reference sequence. Intronic sequence upstream of exon 2 and 500 bp on either end of the intronic sequence downstream of exon 2 are included. **B)** Design of primer/probe pairs used to interrogate splicing of the minigene. Note that codon optimization in exon 3 (underlines) enables these pairs to discriminate the minigene from endogenous PRNP in HEK293 cells.

Using this 6.5 kb minigene as a template, we designed a panel of splice site modifications that we hypothesized would strengthen exon 2 inclusion in the context of human *PRNP* (Figure 3A). These included 1) installation of the consensus strongest^31^ human splice donor and acceptor (“canonical ss”; 6 nucleotide changes required), 2) installation of the mouse *Prnp* exon 2 splice sites (“mouse ss”; 5 changes required), 3) conversion of the donor +5 site from A to G, as this site shows the strongest nucleotide preference of any extended splice site position^26^ (D+5, A>G); 4) conversion of the acceptor -3 site from A to C, to assess whether this single change could mimic the effect of installing the consensus human splice site (A-3, A>C); and 5) conversion of the acceptor -4 site from T to A, to assess whether this single change could mimic the effect of installing the mouse *Prnp* splice site (A-4, T>A).

**Figure 3.**
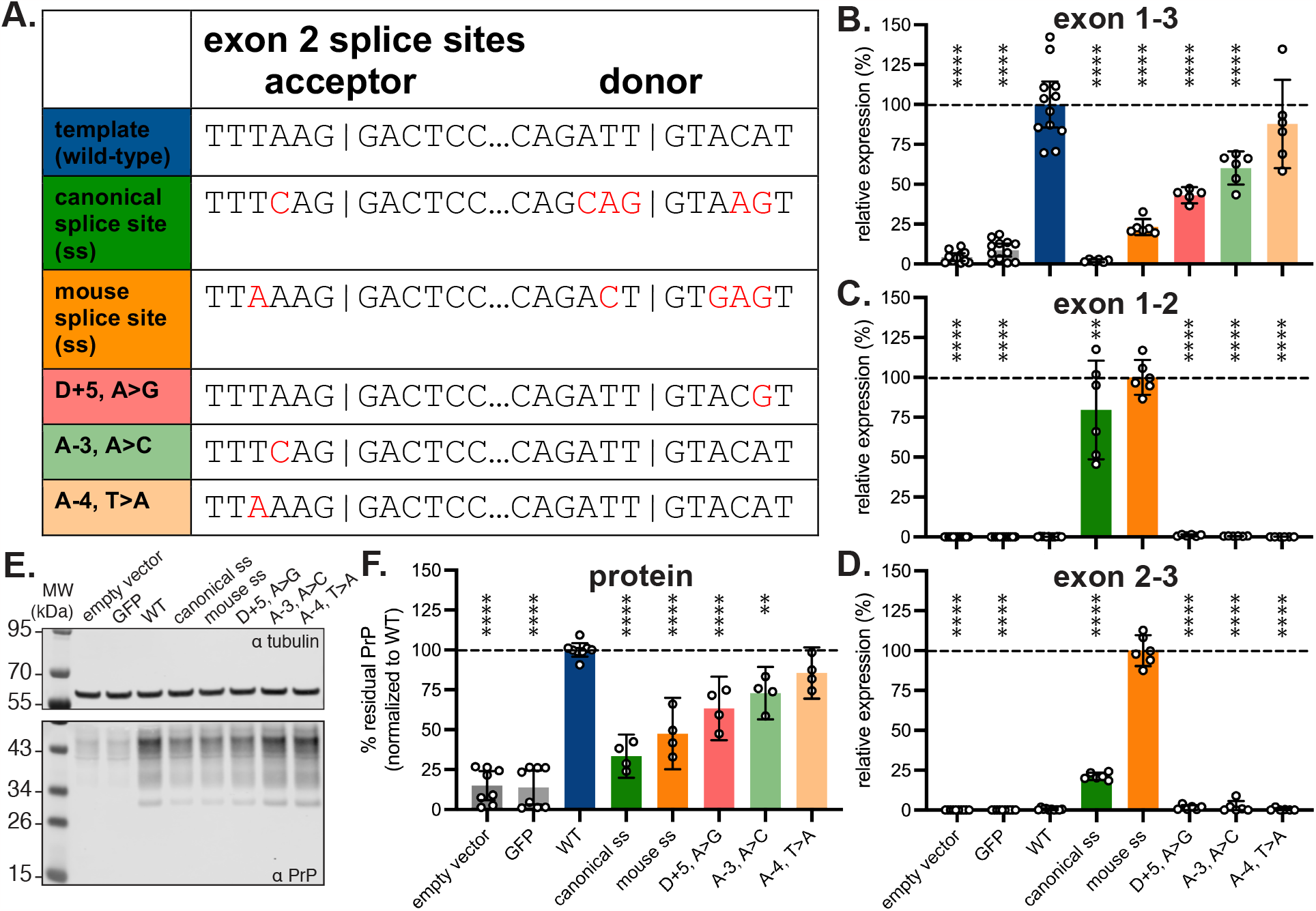
Inclusion of exon 2 lowers PrP expression. **A)** Sequence variants of minigene tested in HEK293 cells. **B-D)** Expression of (B) exon 1-3 (n = 5-12 transfected wells/variant), (C) exon 1-2 (n = 6-12 transfected wells/variant), and (D) exon 2-3 (n = 6-10 transfected wells/variant) junctions in minigene mRNA for each variant transfected into HEK293 cells. Normalized to the template minigene for exons 1-3 and normalized to the highest-expressing variant for exons 1-2 and 2-3. Note that codon optimization in exon 3 enables discrimination from endogenous PRNP. **E)** Immunoblot (POM2 primary antibody^32^ of PrP expression in HEK293 cells transfected with each variant. **F)** Quantification of PrP expression from ≥4 immunoblots per construct, n = 4-8 transfected wells/variant.

Each mutant was separately transfected into HEK293 cells alongside the parent minigene construct and empty vector and GFP transfection controls and analyzed by qPCR. Each primer/probe set (Figure 2B) was designed to amplify only if the targeted exons are adjacent. In keeping with these expectations, the empty vector and GFP controls yielded negligible signal for all primer pairs; trace amplification of exon 1-3 may reflect imperfect allele specificity, as only 2 bases differ from endogenous *PRNP* in the exon 3 codon-optimized primer. The parent minigene yielded PCR product for exon 1-3 but not for exon 1-2 or 2-3, reflecting the baseline exclusion of cryptic exon 2 in a human system.

All 5 splice site mutants appeared to reduce the amount of normally spliced *PRNP* RNA, as measured by the exon 1-3 primer pair, with the change being significant for 4 mutants (Figure 3B). The two mutants yielding the greatest reduction — the canonical ss and mouse ss mutants — showed a corresponding increase in the presence exon 1-2 and 2-3 junctions (Figure 3C-D). For all other constructs, exon 2 remained undetectable, or nearly so, by these primer/probe sets. Note that the results for exons 1-2 and 2-3 are normalized to the highest value obtained for any mutant; 100% does not necessarily mean 100% exon 2 inclusion.

Immunoblots on cell lysates revealed apparent reductions in PrP for all mutants tested (Figure 3E-F). The canonical ss mutant yielded 33% and the mouse ss mutant 48% of the PrP expression level of the parent minigene (Figure 3F). HEK293 cells express endogenous PrP, however, at ∼15% the level achieved by transfection of the parent minigene (Figure 3E-F); adjusting for this floor yielded residual PrP expression of 22% and 38% for the canonical ss and mouse ss mutants, respectively. Across all mutants, PrP levels tracked closely with exon 1-3 qPCR results, with significant reductions for mutants D+5, A>G and A-3, A>C despite the lack of detectable exon 1-2 and 2-3 junctions (Figure 3F, 3C, 3D).

## Discussion

We find that splice site manipulation can modulate the level of PrP in a human cell system, reducing the levels of this disease-causing protein by 78% in the strongest condition tested. For the strongest mutants, which incorporated 5-6 nucleotide changes across the splice donor and acceptor sites, this reduction in protein level was observed in tandem with exon 2 inclusion at the mRNA level. This would be consistent with uORF-mediated translational repression, however, we cannot rule out that nonsense-mediated decay (NMD) may be at work, with the exon 1-2 and 2-3 qPCR simply picking up the small fraction of exon 2-including mRNA that has not yet been degraded. NMD was long held to require 50 bp of distance between the stop codon and the splice donor^33^, versus only 22 bp here, but data from protein-truncating variants in human tissues show this is not a hard-and-fast rule, and that distance from the splice donor is but one of many imperfect predictors of NMD^34^. Still, the evidence for NMD caused by uORFs in human genes is equivocal^18,35^. The single point mutants tested here reduced PrP and normal exon 1-3 splicing without yielding detectable exon 1-2 and 2-3 splicing. Thus, additional mechanisms not foreseen by our initial hypothesis could be operative. One possibility is that these mutants cause inclusion of exon 2 but also retention of a portion of intronic sequence, causing the exon 1-2 and 2-3 qPCR to not amplify, while still resulting in NMD and/or uORF-mediated translational repression.

Our study has several limitations. The battery of splice manipulations that we tested was limited, leaving open the possibility that other splice site changes could yield more dramatic results. As our experiments were limited to human cell culture, in vivo relevance was not demonstrated. Most importantly, the genetic engineering used to establish this proof of concept does not offer a direct path to therapeutic application.

In principle, several therapeutic modalities could be deployed to modulate *PRNP* splicing^36,37^. Antisense oligonucleotides (ASOs) are a well-established modality capable of causing exon inclusion^38^, but may be unlikely to be deployed towards this end: given the desired mechanism of reducing *PRNP* expression, RNAse H1 ASOs are likely to yield greater target suppression than splice-modulating ASOs. Adenine base editors have been successfully deployed to disrupt splice sites^39,40^, however, the single point mutants identified here had relatively modest effects on PrP expression. Instead, small molecule modulation of *PRNP* splicing is the most enticing possibility suggested by our results. PrP-lowering small molecules could have desirable pharmacologic properties, particularly in terms of distribution to deep brain structures less well-reached by oligonucleotides^41^. Attempts to discover small molecules to bind PrP have been unsuccessful^42^, so splicing could offer a new mechanism for small molecule therapies in prion disease. Because *PRNP* does not share the preferred splice site motifs of any known splice-modulating small molecule series^10,12,14^, discovery of a modulator would require a new screening effort.

Despite these limitations, we are encouraged to discover a novel mechanism by which PrP expression can be influenced. PrP’s role in prion disease is uniquely pivotal, as it serves as protein-only pathogen, amplification substrate, and mediator of neuronal neurotoxicity. The therapeutic benefit of PrP lowering has been shown across multiple prion strains^4^, both through genetic reduction and by use of antisense oligonucleotides, and evidence for tolerability is provided by multiple nonhuman species as well as human genetics^43–48^. Given this clarity, PrP and its precursors are disease targets worthy of ongoingly creative angles of attack.

## Methods

### Kozak sequences

Files were retrieved from the Matched Annotation from NCBI and EMBL-EBI (MANE) database (version 1.0) https://ftp.ncbi.nlm.nih.gov/refseq/MANE/MANE_human/release_1.0/ cDNA Transcript sequences from the MANE.GRCh38.v1.0.refseq_rna.fna.gz file were then filtered to only MANE Select transcripts and an 11-bp context surrounding the CDS start were extracted, excluding transcripts where a 11bp CDS context could not be retrieved, as in the case for a leaderless mRNA. To generate a sequence logo, these 11-bp sequences were then super-imposed to align with each other and plotted using ggplot2 and the R package ggseqlogo using the bits method. To generate a histogram, the relative translational efficiencies of each sequence were taken from Noderer et al^27^ and normalized to the most efficient Kozak sequence.

### Comparative genomic analyses

*PRNP* sequences, and multiple alignments thereof, were obtained from UCSC Genome Browser^49^ (accessed September 6, 2023). Kozak sequence strength percentiles were obtained from the rank order among all possible Kozak sequences reported by ref ^27^. GTEx^29^ RNA-seq coverage data were obtained from UCSC Table Browser (accessed November 14, 2023). Exon 1 and 2 in diagrams correspond to the canonical Ensembl transcript ENST00000379440.9.

### Cell culture and transfections

HEK293 cells were maintained in DMEM/F-12 (Gibco, cat no. 11320033) supplemented with 1% Penicillin-streptomycin (Gibco, cat no. 15140163) and 10% FBS (Gibco, cat no. 16000044). For transfection, cells were plated in a 12-well or 96-well plate for protein or RNA analysis, respectively, and were allowed to adhere for 18 hours. Cells were then transfected using Lipofectamine 3000 transfection reagent (Invitrogen, cat no. L3000015) according to the manufacturer protocol. In short, lipofectamine 3000 reagent was diluted in Opti-MEM I reduced serum media (Gibco, cat no. 31985088) for a final mixture containing 3% lipofectamine 3000. In a separate tube, 1 μg (12-well plate) or 0.1 μg (96-well plate) DNA was mixed with 4% P3000 reagent in Opti-MEM. The two tubes were slowly mixed then allowed to incubate at room temperature for 10 minutes before applying the mixture to the cell media. Transfection was incubated on cells for 48 hr before lysing cells.

### Plasmids

Plasmid cloning was performed by Genscript using a modified version of pcDNA3.1(+). The CMV promoter was cloned out of the pcDNA3.1(+) backbone (addgene V790-20) by digesting the vector with NruI and NheI. The human PGK promoter (addgene 82579) was cloned into the backbone, creating pcDNA3.1(+)-hPGK. The minigene was synthesized with codon optimized exon 3, then was ligated into pcDNA3.1(+)-hPGK between NheI and EcoRI.

### Western blot analysis

Following the 48 hr transfection, cells were washed thoroughly with ice-cold PBS then were lysed in 0.2% CHAPS containing cOmplete™, Mini, EDTA-free Protease Inhibitor Cocktail (Sigma, cat no. 4693159001). Protein concentration was determined using a DC protein assay kit (Bio-rad, cat no. 5000112). NuPAGE 4-12%, Bis-Tris, mini protein gels (Invitrogen, cat no. NP0323BOX) were loaded with 10 μg total protein for each sample and run at 180 V in 1x MES buffer (Thermo, cat no. NP0002). Gels were transferred to PVDF membranes using an iBlot 2 device (iBlot™ 2 Transfer Stacks, PVDF, mini, Thermo, cat no. IB24002), 20 V, 7 minutes. Membrane was then cut right under 55 kDa band before blocking with LICOR TBS blocking buffer (LICOR, cat no. 927-60001), 1 hr at room temperature. Primary antibodies were diluted in LICOR TBS blocking buffer + 0.2% Tween-20 (Teknova, cat no. T0710) and incubated at 4°C overnight: α-Tubulin (Invitrogen, cat no. A11126), final 100 ng/μL; POM2 (Millipore, cat no. MABN2298), final 50 ng/μL; 6D11 (BioLegend, cat no. 808001), final 2 μg/μL. Membranes were washed in 1x TBST then incubated in secondary antibody (IRDye® 800CW Goat anti-Mouse IgG, LICOR, cat no. 926-32210) diluted in LICOR TBS blocking buffer + 0.2% Tween-20 and incubated at room temperature for 1 hr. Membranes were again washed with 1x TBST then scanned on a LICOR Odyssey CLx Infrared Imaging System. Blots were analyzed in Fiji^50^.

### qPCR

Following the 48 hr transfection, cells were lysed using the Cells-to-CT 1-step Taqman Kit (Invitrogen, cat no. A25602) using the manufacturer protocol. In short, media was aspirated, each well was washed with 200 μL of ice cold 1x PBS then wash was completely aspirated. Room temperature DNase/Lysis solution (0.5 μL: 50 μL) was added to the cells then plate was put on a shaker for 5 minutes. Finally, 5 μL of room temperature stop solution was added to the cells then plate was put back on the shaker for 2 minutes before moving the plate to ice. RT-PCR samples were prepared using Taqman 1-Step qRT-PCR master mix and Taqman gene expression assays for human *TBP* (Invitrogen, cat no. Hs00427620_m1). Custom primers and probes were ordered from Genscript to quantify the different splice variants (see Table S1 for sequences and Figure 2 for alignment on the minigene sequence). Samples were run on a QuantStudio 7 Flex system (Applied Biosystems) using the following cycling conditions: Reverse transcription (RT) 50°C, 5 min; RT inactivation/initial denaturation 95°C, 20 sec; Amplification 95°C, 3 sec, 60°C, 30 sec, 40 cycles. Each biological sample was run in duplicate and the level of all targets were determined by ΔΔCt whereby results were first normalized to the housekeeping gene *TBP* and then to the wild-type template (exon 1-3) or the mouse ss (exon 1-2 and 2-3), depending on the primer pair used.

### Experimental design and statistical analysis

All data was generated from at least 3 independent transfections, N are as indicated in figure legends. Throughout, all error bars in figures represent 95% confidence intervals. All data were compared with an ordinary one-way ANOVA and Dunnett’s multiple comparison test, with a single pooled variance. P values less than 0.05 were considered nominally significant. In plots, **, p < 0.01 and ****, p < 0.0001. Intron/exon diagrams were plotted in R, qPCR analysis was performed in Google Sheets, and barplots and statistical analyses were performed in Graphpad Prism. Raw data, Prism files, and source code will be made available at https://github.com/ericminikel/cryptic_exon

## AUTHOR CONTRIBUTIONS

Dr. Vallabh had full access to all of the data in the study and takes responsibility for the integrity of the data and the accuracy of the data analysis. Concept and design: SMV, EVM, JEG. Sample analysis: JEG, TLC, MAM. Statistical analysis: JEG. Genomic analysis: EVM, NW, END. Drafting of the manuscript: EVM and SMV. Critical review of the manuscript: all authors. Obtained funding: SMV, NW.

## SUPPLEMENT

Supplementary tables 1-6 provided as a separate Excel file.

**Figure S1.**
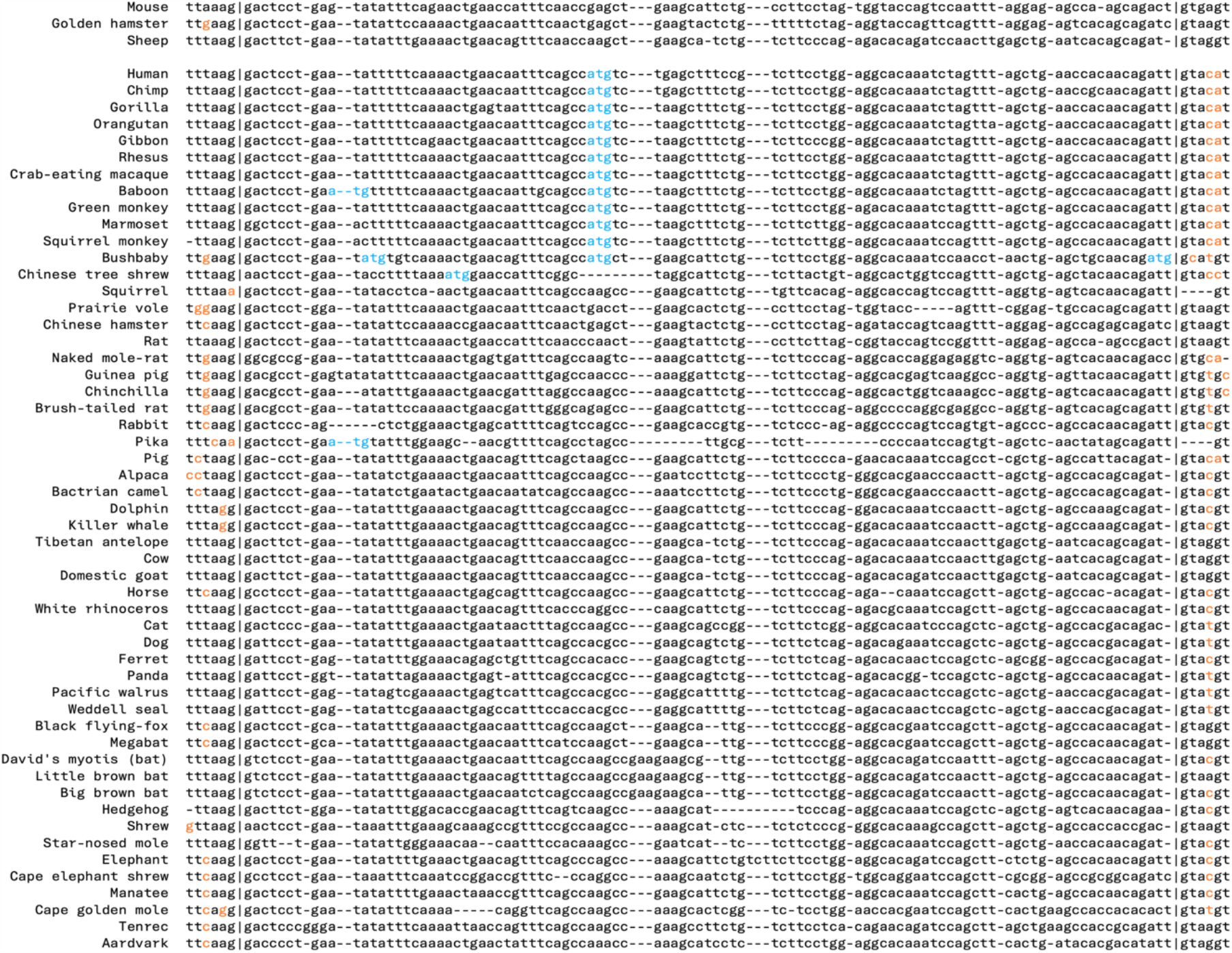
Multiple alignment of PRNP exon 2 orthologous sequence for all available eutherian mammals. As in Figure 1D, but including eutherian mammals without ATGs in exon 2. Lesser Egyptian jerboa is excluded because orthologous sequence was identified for only part of exon 2.

**Figure S2.**
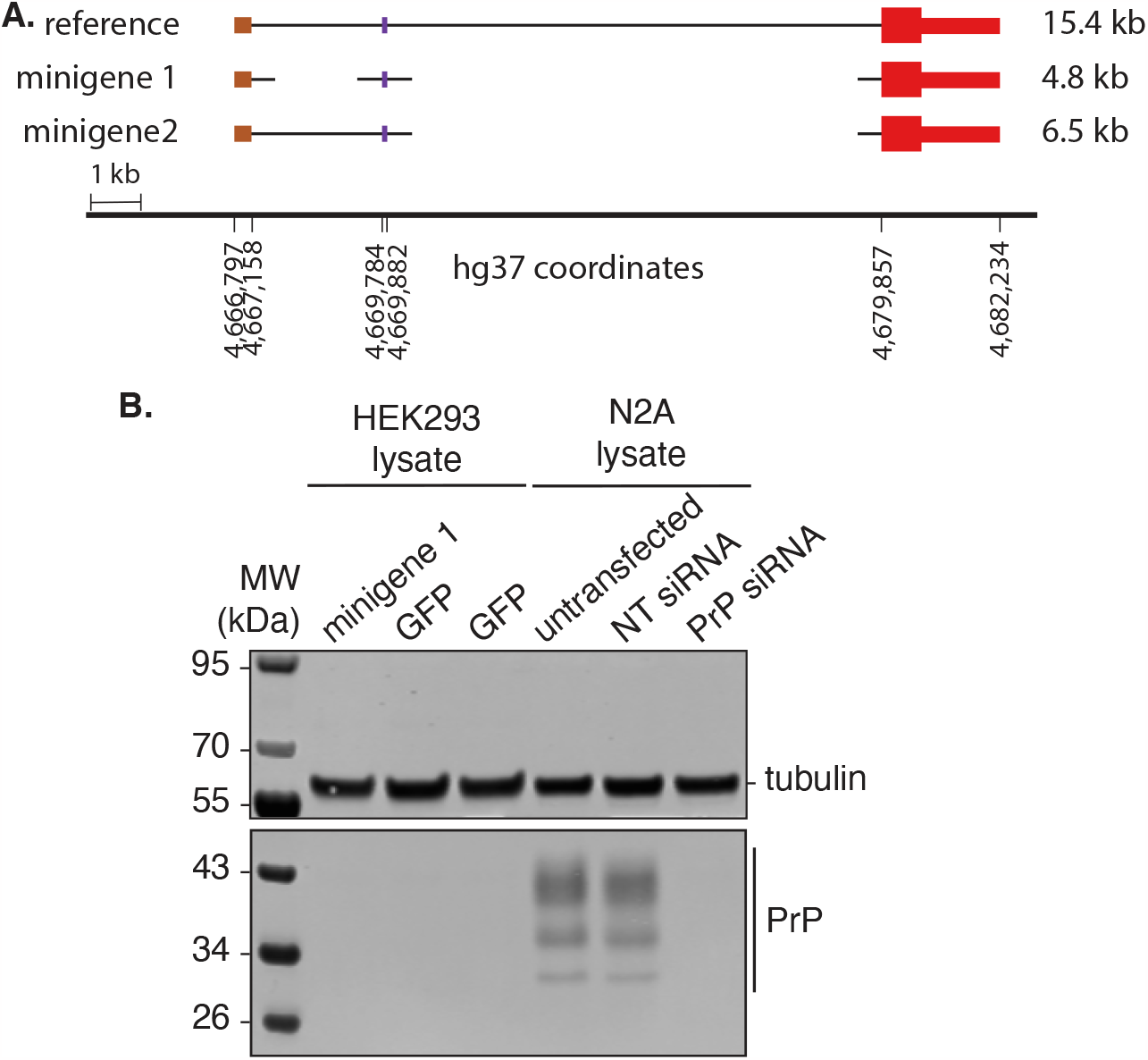
Alternative minigene construct tested in cells. **A)** Comparison of human reference sequence with an alternative “minigene 1” containing only 500 bp at either end of intron 1, and the “minigene 2” used throughout the main text of this manuscript. **B)** Immunoblot failing to detect any expression of minigene 1 in transfected HEK293 cells. Primary antibody: 6D11, see Methods.

